# Screening of the antimicrobial activity of marine sponge extracts from Martinique

**DOI:** 10.1101/2021.02.16.431398

**Authors:** Julie Piron, Jessica Pastour, Niklas Tysklind, Juliette Smith-Ravin, Fabienne Priam

## Abstract

Marine sponges are known for their antimicrobial, antifungal, and cytotoxic activity. In this study, the activity of aqueous and ethanoic extracts of 3 sponges from Martinique were tested on 5 bacterial strains: *Bascillus cereus* (CIP 783), *Echerichia coli* (CIP 54127), *Pseudomonas aeruginosa* (CIP A22), *Staphylococcus aureus* (CIP 67.8) and *Staphylococcus saprophyticus* (CIP 76125). The antimicrobial activity of *Agelas clathrodes, Desmapsamma anchorata*, and *Verongula rigida*, was demonstrated using the disc diffusion method and by determining the minimum inhibitory concentration and the minimum bactericidal concentration. The ethanoic extract of *Agelas clathrodes* had an inhibitory activity specifically on *Staphylococcus aureus* and *Staphylococcus saprophyticus*. No activity was observed for the other extracts. Further chemical analyses will be carried out in order to identify the active molecules of these sponges.

## Introduction

New natural active biomolecules are constantly being sought. In pharmacology, they account for around 70% of the biomolecules used [1], and the potential of those produced by marine organisms is of particular interest given that oceans cover the majority of the earth’s surface. In 2017, seven pharmaceutical products derived from marine substances were approved for clinical use by the FDA [2]. Antibiotics in particular are sought after due to the increasing resistance of many bacterial strains to commercial antibiotics, such as the ‘priority pathogens’ *Pseudomonas aeruginosa* and *Staphylococcus aureus* [3].

This study focuses on the search for marine sponges present in Martinique with antibacterial activity. Marine sponges produce abundant secondary biomolecules and have highly diversified chemical natures [3-6]. The marine ecosystems of Martinique are rich in sponges [7–12], however, they are undervalued [13]. The aim of this study is to increase our knowledge of Martinique’s sponges and their potential antimicrobial activity. One species was found to have high potential antimicrobial activity.

## Materials and Methods

### Sampling and identification

Samples of *Agelas clathrodes, Desmapsamma anchorata*, and *Verongula rigida* were taken from the ‘Fond Boucher’ site (14°39’21.12’’N-61°9’21.22’’W), on the western coast of Martinique (Table 1). Two sampling dives (7/10/2017, 26/11/2018) at 15-20m depth were conducted by Dr. Romain Ferry. Samples were placed in individual plastic bags and immediately stored in a cooler for transport to the University of the West Indies laboratory for analysis. Samples were washed with fresh water and stored at -20°C before extraction.

**Table 1.** Sponge sampling sites

| Species | Site | GPS coordinates |
| --- | --- | --- |
| *Agelas clathrodes* | Fond Boucher | 61° 9'21.22"W 14°39'21.12"N |
| *Desmapsamma anchorata* | Fond Boucher | 61° 9'21.22"W 14°39'21.12"N |
| *Verongula rigida* | Fond Boucher | 61° 9'26.55"W 14°39'23.67"N |

The sponge skeletons were studied based on spicules and longitudinal and transverse sectioning of the tissue. The spicules were removed by freeze-drying 5mg samples which were then bleach-washed 3 times. Examination was carried out using an optical microscope equipped with a ZEISS camera. Photographs were analysed using the ZEN2012 software.

### Preparation of sponge extracts

Extracts were prepared by maceration in one of two solvents: distilled water and 100% ethanol. After freeze-drying, 2g samples were cut into small pieces, crushed in a mortar, and transferred to a 15ml falcon tube before 10ml of solvent was added. After repeated manual turning and swirling, the tubes were agitated for 24 hrs at room temperature. The extracts were then filtered through standard filter paper. This process was repeated 3 times before the extracts were pooled and stored at -20°C before drying.

The ethanoic extracts (E) were dried in a rotary evaporator at 45°C. Two rounds of 1mL of solvent were added to the rotary flask, and the mixture was then passed through the ultrasonic bath to recover the extracts, before transferring into glass vials. Solvent residues were evaporated from the vials under a fume hood and the dry extract was stored at -20°C prior to testing.

The aqueous extracts (A) were freeze-dried and stored at -20°C.

## Cultivation of bacterial strains

Inoculums of 5 bacterial strains were cultivated in nutrient agar at 30°C or 37°C. According to the Article R4421-3 of Decree n°2008-244 of 7 March 208-art. (V), from the French law, *Bascillus cereus* (CIP 78.3), *Staphylococcus saprophyticus* (CIP 76125T), and *Echerichia coli* (CIP 54.127) are classed in group 1, non-pathogenic. Pse*udomonas aeruginosa* (CIP A22), and *Staphylococcus aureus* (CIP 67.8) classed in group 2, pathogenic.

## Antimicrobial test

Antimicrobial activity is tested on Mueller Hinton agar using the disc diffusion method [14]. Pure inoculum cultures, placed in petri dishes and incubated for 18h on Nutrient Agar, were inoculated at a concentration of 10^7^UCF/mL. Sterile 6mm diameter discs were impregnated with 20µL of sponge extract (at 500µg/mL) and dried for 15min. The discs were stored at 4°C before being placed on the inoculated agar and the petri dishes were incubated for 24 hrs at 30°C or 37°C. The inhibition diameter (IØ) was then measured. Tests were carried out in triplicates for each species. Ampicillin (10µg, lot 7B5479) was used as positive control for *S. saprophyticus, S. aureus*, and *E*.*coli*; chloramphenicol (30µg, lot 5C5220) was the positive control for *B. cereus;* and fosfomycin (50µg, lot 4L5252) was the positive control for *P. aeruginosa*. Discs impregnated with 20µL of solvent (H2O and EtOH) were used as negative controls.

An inhibition diameter (IØ) greater than 9mm around the disk indicates positive antimicrobial activity [15].

## Determining minimum inhibition concentration (MIC)

MIC is defined as the minimum concentration that inhibits the visible growth of the bacteria tested after 24 hrs. MIC was determined by the method of successive microdilution in a liquid medium modified from Majali *et al*., [14] for the ethanoic extract of *A. clathordes*. The initial sponge extract solutions (1mg/mL) were diluted successively 11 times at 50% with distilled water, and the resulting dilutions distributed on a 96-well plate. The plates were then incubated for 24 hrs at 30°C or 37°C before the optical density was measured on a plate reader at 600nm. The concentration of extract used ranged from 0.488 to 1000µg/mL.

## Determining minimal bactericidal concentration (MBC)

MBC is defines as the minimum concentration necessary to kill bacteria. MBC was also determined for the ethanoic extract of *A. clathrodes* by counting the surviving bacteria in tubes without visible growth using a method modified from [14, 16]. PCA plates were seeded with a strip of a 10µL extract sample using a calibrated seeder and incubated for 24 hrs at 30°C or 37°C. After incubation, the colonies of each strip were counted. Only strips with 30-300 colonies were counted.

If MBC=MIC, the extract was bactericidal. If MBC>MIC, the extract was bacteriostatic, i.e. the extract inhibited bacteria proliferation without causing death [14].

## Results and Discussion

### Antimicrobial activity

Table 2 shows the antimicrobial activity for the 5 non-pathogenic strains. Only the ethanoic extract of *A. clathrodes* shows a low inhibitory effect, on *S. aureus* (IØ: 10.66mm) and *S. saprophyticus* (IØ: 9.5mm). The inhibition diameters obtained for the extract are lower than those obtained for the synthetic antibiotic ampicillin on *S. aureus* (IØ: 40mm) and on *S. saprophyticus* (IØ: 46mm). These results confirm that Gram+ bacteria are more sensitive to extracts than Gram-bacteria due to higher membrane permeability [17].

**Table 2:** Inhibition diameter (mm) of the aqueous and ethanoic extracts of the different species on the 5 strains.

|  | Extract | Strains |  |  |  |  |
| --- | --- | --- | --- | --- | --- | --- |
|  |  | Gram + |  |  | Gram - |  |
|  |  | *S.aureus* | *B.cereus* | *S.saprophyticus* | *E.coli* | *P.aeruginosa* |
| Ampicilline |  | 40 | - | 46 | 26 | - |
| Chloramphénicol |  | - | 25 | - | - | - |
| Fosfomycine |  | - | - | - | - | 17.5 |
| Control - 1 | A | R | R | R | R | R |
| Control - 2 | E | R | R | R | R | R |
| *A.clathrodes* | A | R | R | R | R | R |
|  | E | 10.66 | R | 9.5 | R | R |
| *D.anchorata* | A | R | R | R | R | R |
|  | E | R | R | R | R | R |
| *V.rigida* | A | R | R | R | R | R |
|  | E | R | R | R | R | R |

A:aqueous extract. E: ethanoic extract. R: Resistant. -: not used for this strain.

In similar tests carried out with *Agelas sventres*, such strain-specific antimicrobial activity on staphylococcus strains was not observed. Antimicrobial activity has been observed on: E.coli (ATCC 25922) for methanoic extracts and n-Hexane, on *S. aureaus* (ATCC 25923) with chloroform and hexane extracts, and on *C. albicans* (ATCC 10231) with chloroform extracts [18]. The specificity of antimicrobial activity on *Staphylococcus* observed with *A. clathrodes* probably indicates a difference in the composition of active biomolecules, due either to the species or to the solvent. A test carried out using a crude methanoic extract of the species *Agelas sp*. showed a significant effect on the same strain of *S. aureus* (ATC 25923), but again the extract inhibitory activity were not specific to *Staphylococcus*, and would be due to the presence of two types of molecules, agelasine B and D [17]. Alcohol extracts therefore seem to have a greater effect on staphylococci than on other strains. Alcohol probably enables better extraction of alkaloid-type biomolecules, and marine sponge alkaloids are known for their antibacterial effects, on *S. aureus* particularly. For example, bromo-pyrrole alkaloids extracted from *Agelas dispar* have moderate antimicrobial activity on the Gram + strains *B. subtilis* and *S. aureus* [19]. Similar biomolecules are thus possibly present in the *A. clathrodes* extract. The absence of effect on these same strains with the aqueous extract confirms the need for an alcoholic solvent for alkaloids biomolecules, which seems to be specific to the *Agelas* genus, whence the absence of effect from the *D. anchorata* and *V. rigida* extracts. As for *Agelas dilatata*, pyrrole-imidazoles have been tested on two pathogenic strains of *P. aeruginosa* (ATCC 27853 and PAO1) and showed moderate to strong activity. Bromoageliferin, an isolated molecule, showed significant activity on *P. aeruginosa* strain ATCC 27853 in particular [20]. This indicates the absence of this molecule in *A. clathrodes*.

The negative results observed for *D. anchorata* on all strains were expected, extracts having previously had no antimicrobial effect on *S. aureus, E. coli*, and *P. aeruginosa*. [21]. *D. anchorata* served in this study as a negative control, and our results confirm those previously obtained.

For *V. rigida*, already known for its antiparasitic [17,22], or antidepressant properties [23], our results show the species has no antibacterial properties.

### MIC and MBC

In order to analyse the observed antimicrobial activity, we determined the MIC and the MBC of the ethanoic extract of *A. clathrodes*. MIC is defined as the minimum concentration that inhibits the visible growth of the bacteria tested after 24 hrs, while the MBC is defines as the minimum concentration necessary to kill bacteria. MIC of *A. clathrodes* extract was 15.625 µg/mL for *S. aureus* and 7.813 µg/mL for *S. saprophyticus* (Table 3). Although the effect on *S. saprophyticus* was lower according to the antibiogram, the MIC results show greater activity on *S. saprophyticus* at low concentration. The results were similar for the MBC, 31.250 µg/mL for *S. aureus* and 15.625 µg/mL for *S. saprophyticus* (Table 3). MBC/MIC ratios of 2 indicate that *A. clathrodes* ethanoic extracts have bacteriostatic effects. The extracted biomolecules therefore have an inhibitory effect on the growth of the *S. aureus* and *S. saprophyticus* strains.

**Table 3:**
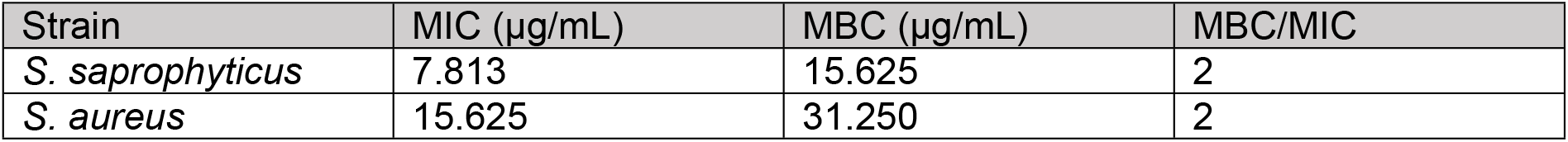
MIC and MBC for *Agelas clathrodes* extrracts.

| Strain | MIC (µg/mL) | MBC (µg/mL) | MBC/MIC |
| --- | --- | --- | --- |
| *S. saprophyticus* | 7.813 | 15.625 | 2 |
| *S. aureus* | 15.625 | 31.250 | 2 |

With the quality of the extractions carried out and the quantity of extracts obtained after drying, a higher MIC for *S. aureus* indicates a lower concentration, not absence, of active biomolecules. In fact, a higher MIC has already been observed for an alcoholic extract of sponge *Stylissa massa* on *E*.*coli*, [24]. The bacteriostatic effect of *A. clathrodes* extract may, once again, indicate the presence of biomolecules different from those known on other species of the *Agelas* genus.

Future complementary analyses, including cytotoxic tests and LC-MS analyses of these extracts, will allow further exploration of this hypothesis.

## Conclusion

Preliminary tests of raw extracts of three marine sponges from Martinique have allowed us to determine their potential antimicrobial activity. *Agelas clathrodes* has an observed antimicrobial activity on staphylococcus strains, a pathogenic bacterium with increasing resistance to antibiotics. Its exploitation could provide a potential alternative to synthetic antibiotics. The presence of species of pharmacological interest, such as *A. clathrodes*, is a key element in the development of the natural heritage of Martinique. This study is the first of its kind for Martinique, and its results will be further investigated, through the chemical screening of extracts and the analysis of chemical compounds, in order to determine the differences in the composition of extracts according to the solvent and thus to identify the active biomolecules. This study is also the first of its kind for *Verongula rigida*, which does not show any antimicrobial activity on the strains tested. Screening could be broadened to strains not included in our study.

Further analysis could also determine whether the biomolecules in these extracts are originate from the sponges or from symbiotic organisms such as fungi. [25,26] or bacteria [27].

## Acknowledgements

The authors thank Dr. Ferry for his involvement in the sampling and identification of the sponges. They also thank Dr. Stéphanie Morin for her help in conducting antibiograms.

